## supplemental materials for "The nanoscale organization of the Nipah virus fusion protein informs new membrane fusion mechanisms"

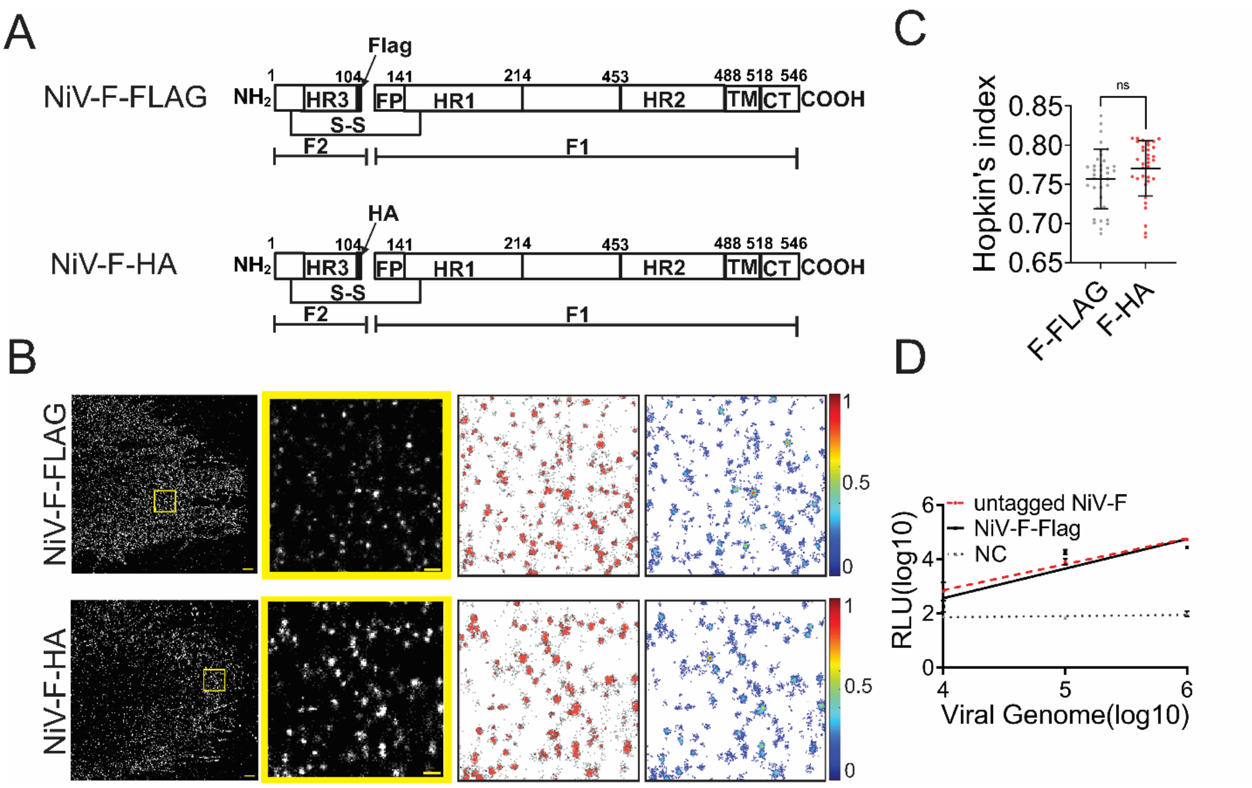


***Fig. S1 (related to Fig. 1). A comparison of clusters formed by NiV-F-FLAG and NiV-F-HA on PK13 cells.*** *(A) A diagram of NiV-F-FLAG (Top) and NiV-F-HA (Bottom) constructs. Both tags were inserted after amino acid 104 of NiV-F. (B) First column**:** Cross-section (Δz = 600 nm) of SMLM images of NiV-F-FLAG (top) and NiV-F-HA (bottom) in PK13 cells. Scale bar: 1 μm. Second column: The yellow boxed region is enlarged to show the detailed distribution pattern. Scale bar: 0.2 μm. Third column: Cluster maps of the enlarged regions. Fourth column: Localization density maps show the normalized relative density of the enlarged regions. (C) Hopkin's index of the NiV-F-FLAG and NiV-F-HA localizations in PK13 cells. n = 36 and 34. Sample size n is the number of total regions from 4 cells. Bars represent mean ± SD. P value was obtained using the Mann-Whitney test. ns: p > 0.05; * p<0.01; ** p<0.001; *** p<0.0001. (D) The entry of VSV/NiV pseudoviruses expressing NiV-G with untagged NiV-F (red), NiV-F-FLAG (black), and pcDNA 3 vector (NC; gray) in Vero cells. Data shown are mean ± SEM from one representative experiment (of three).*

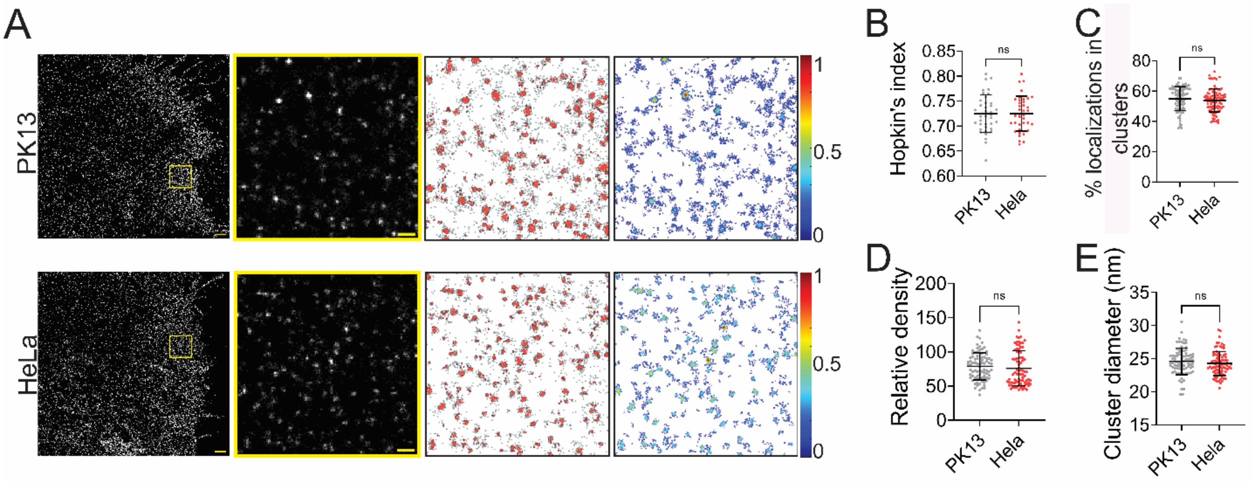


***Fig. S2 (related to Fig. 1). The nanoscale organization of NiV-F is similar in PK13 and HeLa cells.*** *(A) First column: Cross-section (Δz = 600 nm) of SMLM images of NiV-F in PK13 (top) and HeLa (bottom) cells. Scale bar: 1 µm. Second column: The yellow boxed region is enlarged to show the detailed distribution pattern. Scale bar: 0.2 µm. Third column: Cluster maps of the localizations in the enlarged region. Fourth column: localization density maps show the normalized relative density of the enlarged regions. (B-E) Quantitative analyses of clustering of NiV-F in PK13 and HeLa: (B) Hopkin's index, n = 38 and 38; (C) Percentage of localizations in clusters, n = 84 and 90; (D) Relative density, n = 84 and 100; (E) Average cluster diameters, n = 84 and 84. Bars represent mean ± SD. Sample size n is the number of total regions from 4-10 cells. p value was obtained using the Mann-Whitney test. ns: p > 0.05; * p<0.01; ** p<0.001; *** p<0.0001.*

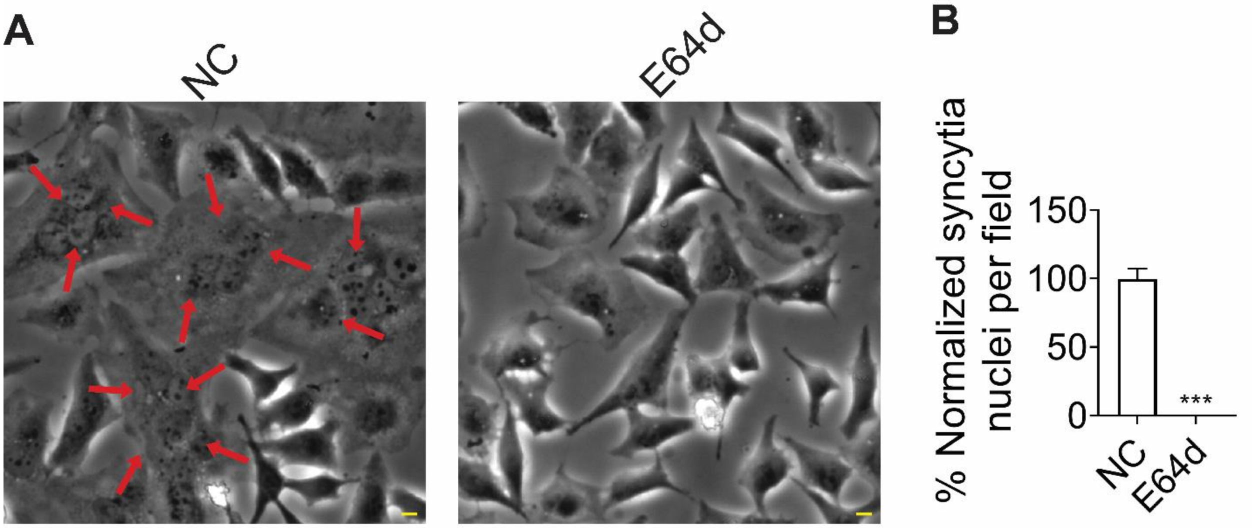


***Fig. S3 (related to Fig. 2). E64d treatment inhibits cell-cell fusion induced by NiV-F and -G.*** *(A) Representative images of cell-cell fusion induced by NiV-G and NiV-F without (NC) or with (E64d) the treatment of 20 μM E64d in HeLa cells. Cells were co-transfected by expression plasmids coding for NiV-G and NiV-F, at 3 hrs post-transfection, 20 μM E64d or the same volume of solvent methanol was added to cells. Cells were fixed at 18 hrs post-transfection and observed under 20 x magnification. Scale bar = 10 μm. (B) Cell-cell fusion levels normalized to that of the untreated HeLa cells (NC). Five fields per experiment were counted from three independent experiments. Bars represent mean ± SEM. *** p = 0.0002. p value was obtained using Welch's t-test.*

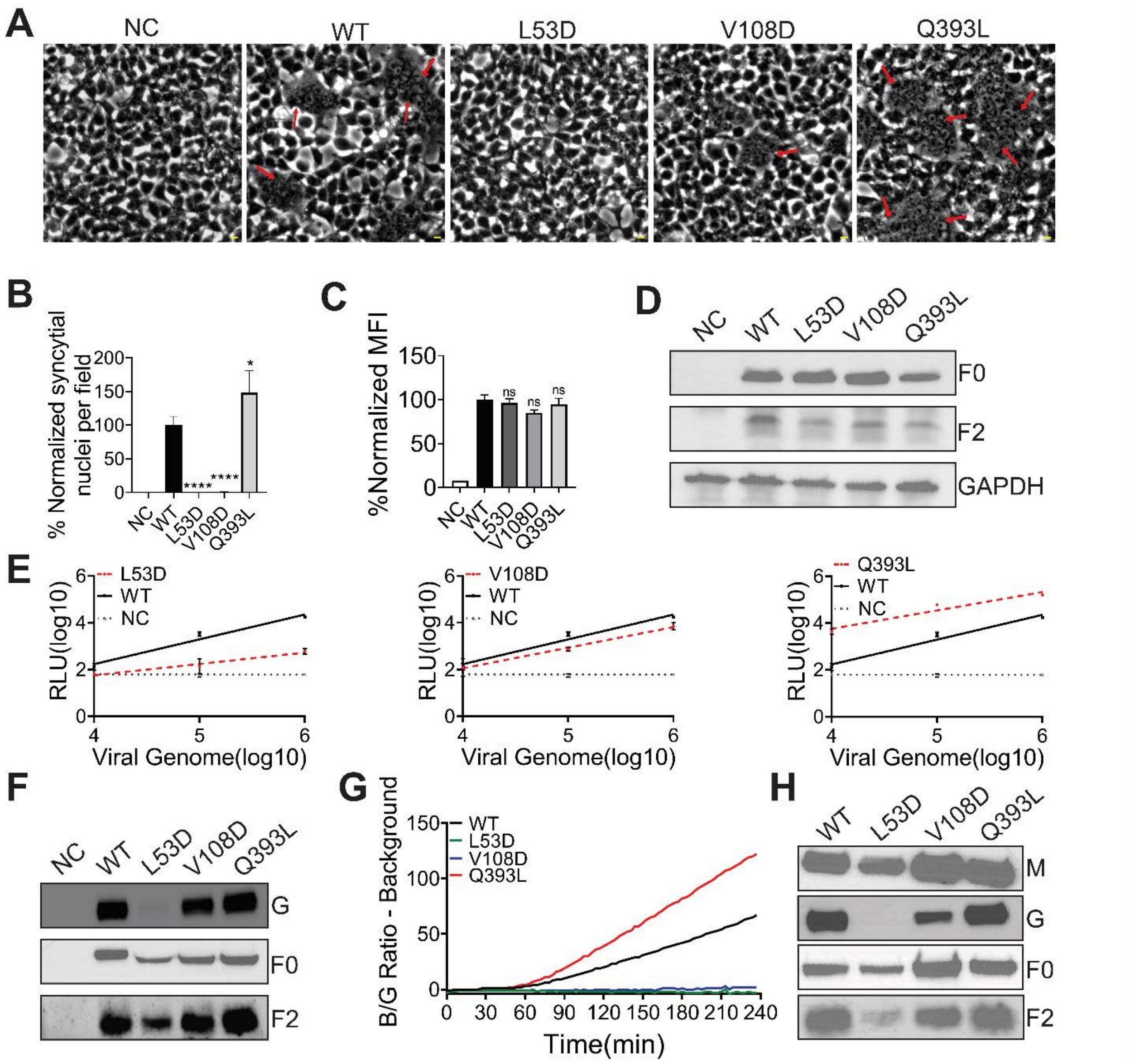


***Fig. S4. (related to Fig. 3). The fusion ability, expression levels, and processing of FLAG-tagged NiV-F and hexameric mutants.*** *(A) Representative images of 293T cell-cell fusion induced by NiV-G and NiV-F-WT, L53D, V108D, or Q393L. 293T cells were co-transfected with plasmids coding for NiV-G and empty vector (NC) or NiV-F constructs. Cells were fixed at 18 hrs post-transfection. Arrows point to syncytia. Scale bar: 10 µm. (B) Relative levels of 293T cell-cell fusion in (A). Five fields per experiment were counted from three independent experiments. Data are presented as mean ± SEM. (C) The cell surface expression levels of NiV-F-WT, L53D, V108D, and Q393L on 293T cells were measured by flow cytometry. Mean fluorescence Intensity (MFI) values were calculated by FlowJo and were normalized to that of F-WT. Data are presented as mean ± SEM of three independent experiments. Statistical significance was determined by the unpaired t-test with Welch's correction (*P<0.05, **P<0.01, ***P<0.001, ns: not significant). Values were compared to that of the NiV-F-WT. (D) NiV-F processing of F-WT, L53D, V108D, Q393L in 293T cells. 293T cells were transfected by F-WT and the mutants. The cell lysates were analyzed on SDS-PAGE followed by western blotting after 28hrs post-*

*transfection. F0 and F2 were probed by M2 monoclonal mouse anti-FLAG antibody. GAPDH was probed by monoclonal mouse anti-GAPDH. (E) Relative entry levels of VSV/NiV pseudovirions expressing NiV-G-HA and NiV-F-FLAG (WT; solid black line) or FLAG-tagged NiV-F-L53D (L53D), V108D (V108D), or Q393L (Q393L; dotted red line). The negative control (NC), the recombinant VSV pseudoviruses without glycoproteins, is shown as a dotted gray line. The relative light units (RLU) of the lysates of infected Vero cells were quantified 18–24 h post-infection and plotted against the number of viral genomes/ml over 3 logs of viral input. Data shown are mean ± SEM from one representative experiment (of three). (F) The result of a representative western blot analysis of VSV/NiV pseudovirions. $4x10^8$ copies VSV/NiV pseudovirions were separated by a denaturing 10% SDS–PAGE and probed against NiV-G-HA (rabbit anti-HA) and NiV-F-FLAG (mouse anti-Flag). (G)The VLPs expressing NiV-M-Bla, G-HA, and the FLAG-tagged WT or mutant NiV-F were allowed to bind to the target HEK293T cells loaded with CCF2-AM dye at 4°C. The Blue/Green (B/G) ratio was measured at 37°C for 4 hrs at a 3-min-interval. The average background was subtracted, and the results were normalized to the maximal B/G ratio of WT VLPs. Results from one representative experiment (of three) are shown. (H) The result of a representative Western blot analysis of NiV VLPs. Equal volumes of VLPs were separated by a denaturing 10% SDS–PAGE and probed against NiV-G (rabbit anti-HA), NiV-F (mouse anti-Flag), and NiV-M (mouse anti- β-lactamase).*

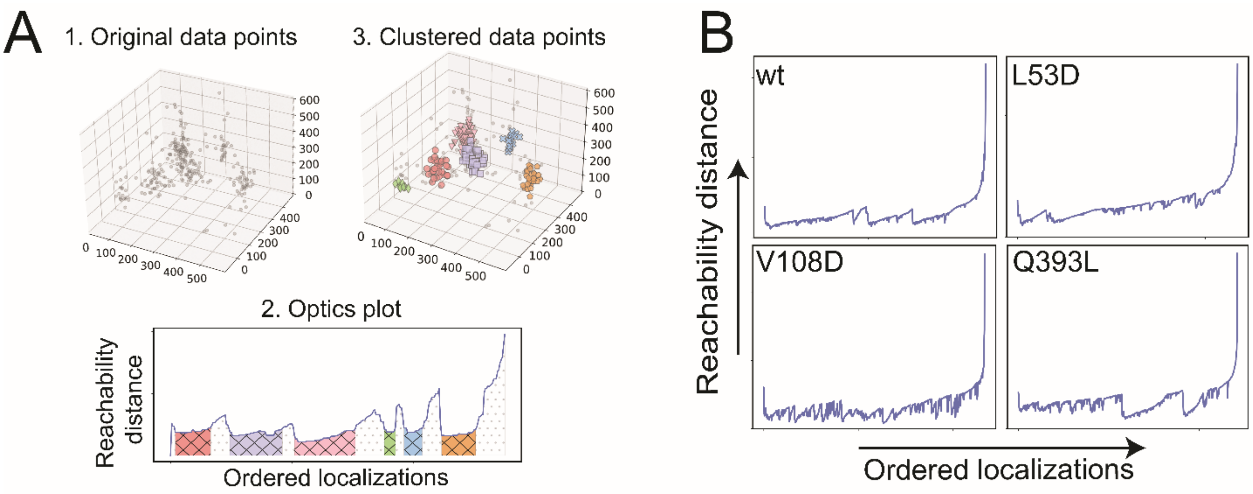


***Fig. S5. (related to Fig. 4) The OPTICS algorithm identifies the clusters of NiV-F-WT and constructs on 3D VLPs.*** *(A) The OPTICS plots for a simulated 3D SMLM dataset containing clusters of different sizes and densities. (B) The OPTICS plots of the localizations of NiV-F constructs on individual VLPs. The OPTICS plot shows the correlation between the ordered localization sequence (x-axis) and the reachability distance (y-axis).*

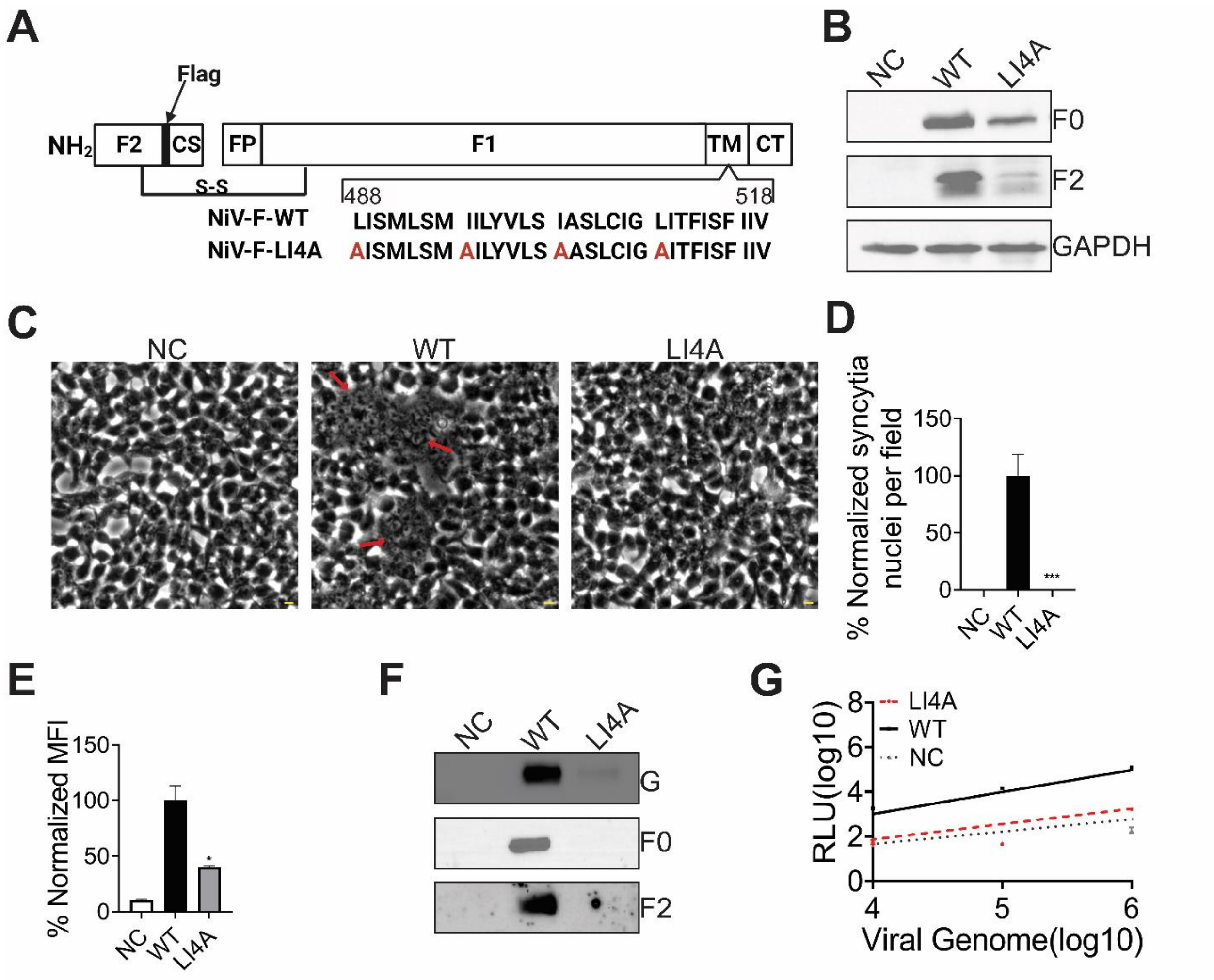


**Fig. S6. (Related to Fig. 5). The NiV-F LI4A does not induce cell-cell fusion.** *(A) A diagram of NiV-F-LI4A (LI4A) mutant that carries alanine mutations at the LI zipper (B) The processing of NiV-F-WT and F-LI4A in 293T cells. Expression plasmids of an empty vector (NC), NiV-F-WT (WT), and NiV-F-LI4A (LI4A) were transfected into 293T cells. At 28 hrs posttransfection, cell lysates were collected and loaded on a 10% polyachrylamide gel for SDS-PAGE. The $F_0$ and $F_2$ were detected using a mouse anti-FLAG antibody and a goat anti-mouse HRP. The GAPDH is a loading control. (C) Representative images of 293T cell-cell fusion induced by WT and LI4A. 293T cells were co-transfected using plasmids coding for NiV-G and an empty vector (NC), NiV-F-WT (WT), or NiV-F-LI4A. Cells were fixed at 18 hrs post-transfection. Arrows point to syncytia. Scale bar: 10 µm. (D) Relative cell-cell fusion levels in (C). Five fields per experiment were counted from three independent experiments. Bars represent mean ± SEM. (E) The cell surface expression levels of WT and LI4A on 293T cells were measured by flow cytometry. Mean fluorescence Intensity (MFI) values were calculated by FlowJo and were normalized to WT. Bars are presented as mean ± SEM of n=3 independent experiments. Statistical significance was determined by the unpaired t-test with Welch's correction (*P<0.05, **P<0.01, ***P<0.001). Values were compared to that of WT. (F) The result of a representative Western blot analysis of VSV/NiV pseudovirions. $4x10^8$ copies of VSV/NiV pseudotyped virions were separated by*

*a denaturing 10% SDS–PAGE and probed against NiV-G (rabbit anti-HA) and NiV-F (mouse anti-Flag). (G) Relative entry levels of VSV/NiV pseudovirions expressing NiV-G-HA and NiV-F-FLAG (solid black line) or LI4A (dotted red line). The negative control (NC), the recombinant VSV pseudoviruses without glycoproteins, is shown as a dotted gray line. The relative light units (RLU) of lysates of infected Vero cells were quantified 18–24 hrs post-infection and plotted against the number of viral genomes/ml over 3 logs of viral input. Data shown are averages ± SEM from one representative experiment (of three).*

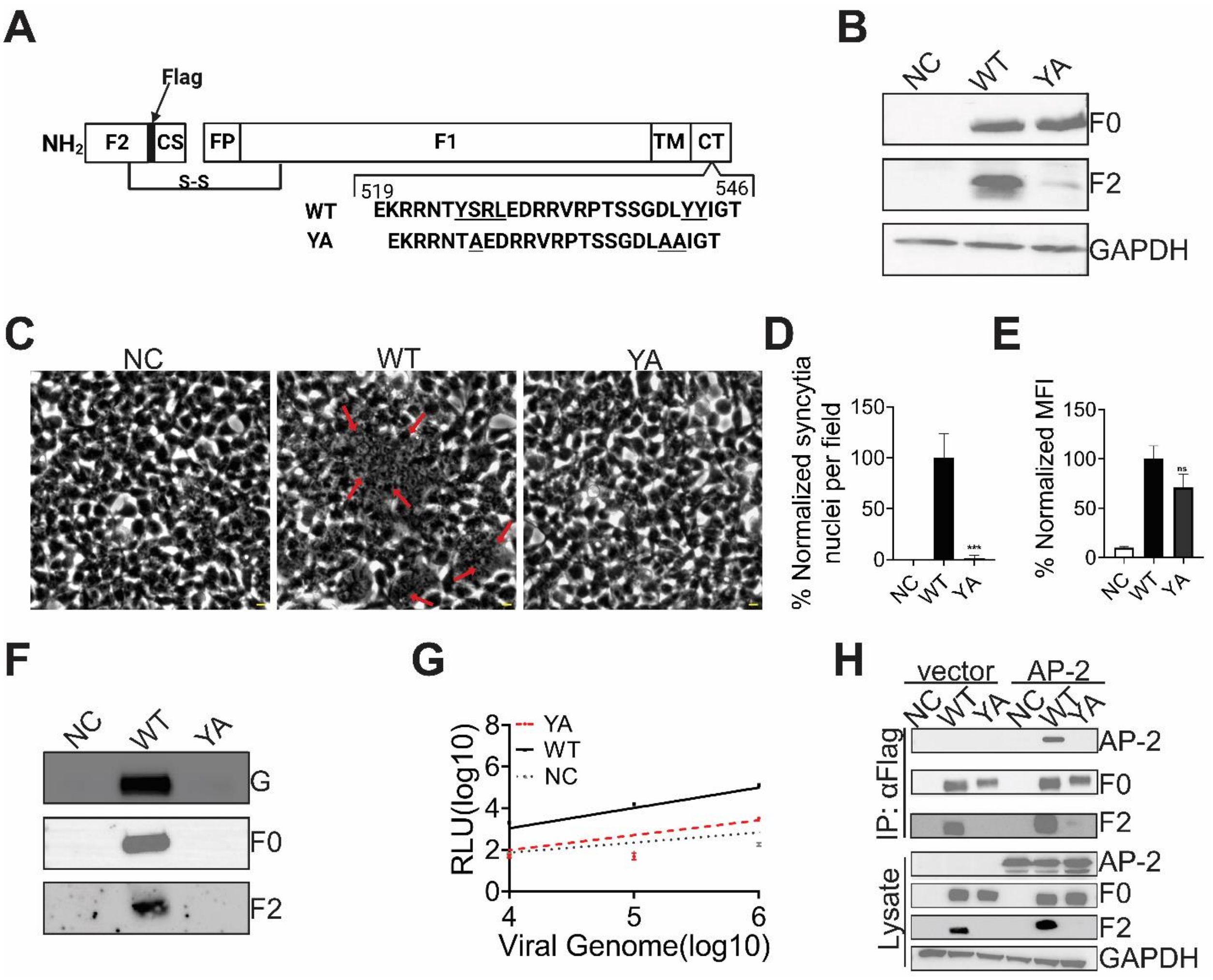


***Fig. S7 (related to Fig. 6). The FLAG-tagged NiV-F-YA mutant inhibits NiV-F cleavage, cell-cell fusion, and the NiV-F-AP-2 interaction in 293T cells.*** *(A) A diagram of NiV-F-YA (YA) mutant which carries alanine mutations of a tyrosine residue at the endosomal sorting signal YSRL and two additional tyrosine residues at the cytoplasmic tail. (B) The processing of NiV-F-WT and F-YA in 293T cells. Expression plasmids of an empty vector (NC), NiV-F-WT (WT), and NiV-F-YA (YA) were transfected into 293T cells. At 28 hrs posttransfection, cell lysates were collected and loaded on a 10% polyachrylamide gel for SDS-PAGE. The $F_0$ and $F_2$ were detected using a mouse anti-FLAG antibody and a goat anti-mouse HRP. The GAPDH is a loading control. (C) Representative images of 293T cell-cell fusion induced by WT and YA. 293T cells were cotransfected using plasmids coding for NiV-G and an empty vector (NC), NiV-F-WT (WT), or NiV-F-YA (YA). Cells were fixed at 18 hrs post-transfection. Arrows point to syncytia. Scale bar: 10 μm. (D) Relative cell-cell fusion levels in (C). Five fields per experiment were counted from three independent experiments. Bars represent mean ± SEM. (E) The cell surface expression levels of WT and YA on 293T cells were measured by flow cytometry. Mean fluorescence Intensity (MFI) values were calculated by FlowJo and were normalized to WT. Bars are presented as mean ± SEM of n=3 independent experiments. Statistical significance was determined by the unpaired t-test with Welch's correction (*P<0.05, **P<0.01, ***P<0.001). Values were compared to that of WT. (F) The result of a representative Western blot analysis of VSV/NiV pseudovirions. $4x10^8$ copies of NiV/VSV pseudovirions were separated by a*

*denaturing 10% SDS–PAGE and probed against NiV-G (rabbit anti-HA) and NiV-F (mouse anti-Flag). (G) Relative entry levels of VSV pseudovirions containing NiV-G-HA and the FLAG-tagged-NiV-F (solid black line) or YA (dotted red line). The negative control (NC), the recombinant VSV pseudoviruses without glycoproteins (VSV/pcDNA3), is shown as a dotted gray line. The relative light units (RLU) of lysates of infected Vero cells were quantified 18–24 h post-infection and plotted against the number of viral genomes/ml over 3 logs of viral input. Data shown are mean ± SEM from one representative experiment (of three) are shown. (H) 293T cells were transfected with the indicated combinations of NiV-F-WT, YA, and AP-2. At 48 hpt, cells were lysed and NiV-F constructs were immunopreciptated by using μMACS. Anti-FLAG magnetic beads were used. Total cell lysate (Lysate) and immunoprecipiated proteins (IP:αFLAG) proteins were separated by 10% SDS-PAGE and immunoblotted with mouse α-FLAG (F detection) or rabbit α-mcherry (AP-2 detection). Proteins were detected using HRP-conjugated secondary antibodies.*
